# SAFE-LD: A novel method for the estimation of linkage disequilibrium from summary statistics

**DOI:** 10.1101/2025.09.29.679154

**Authors:** Giulia Elizabeth de Sanctis, Sodbo Sharapov, Davide Bolognini, Francesca Ieva, Nicole Soranzo, Emanuele Di Angelantonio, Claudia Giambartolomei, Nicola Pirastu

**Author notes:** These authors contributed equally to this work (Claudia Giambartolomei, Nicola Pirastu).

## Abstract

Genome-wide association studies (GWAS) have greatly advanced our understanding of the genetic architecture of complex traits. Downstream analyses of GWAS summary statistics require accurate in-sample LD, the variant correlations in the same individuals used for the GWAS, as even small discrepancies can propagate into substantial error. In practice, privacy and consent restrictions prevent sharing of individual-level genotypes, forcing researchers either to rely on external reference panels, which reduce accuracy and power, or to store and distribute massive precomputed LD matrices that are inflexible and difficult to analyze. Here we introduce **SAFE-LD** (*Shrinkage and Anonymisation Framework for LD Estimation*), a novel method that produces pseudo-genotypes designed to reproduce the exact in-sample LD of a cohort, while discarding all individual-level genetic content. SAFELD surrogates can be stored in VCF/PGEN formats and used seamlessly with standard pipelines, providing LD estimates indistinguishable from the originals but free from privacy concerns. Using extensive simulations on UK Biobank data, we show that SAFE-LD is robust across genomic regions and population sizes. Notably, SAFE-LD achieves finemapping accuracy on par with internal LD, and significantly outperforms external LD even under best-case conditions with cohort-matched reference panels. We further extend this framework to existing GWAS summary statistics through **SAFE-LDss**, which exploits existing published summary statistics where numerous traits have been analyzed on the same samples. SAFE-LD offers a scalable, privacy-preserving, and highly accurate alternative to traditional LD estimation, enabling easy sharing and seamless utilization with standard tools. By storing compact pseudo-genotypes instead of massive precomputed LD matrices, it also provides a highly efficient solution in terms of disk space and data management, while safeguarding participant privacy and supporting precise fine-mapping.

## 1 Introduction

Genome-wide association studies (GWAS) have revolutionized human genetics by enabling the discovery of hundreds of novel genetic associations and supporting a wide range of downstream analyses (Visscher PM [2017], A. [2013]). In addition to gene discovery, many analyses rely critically on a precise estimation of linkage disequilibrium (LD), which captures the correlation structure among single-nucleotide polymorphisms (SNPs). LD information is also central to widely used approaches such as fine-mapping, genetic correlation analysis (Bulik-Sullivan [2015]), polygenic risk score construction (Choi [2020]), and Mendelian randomization (Davey Smith [2014]). In this work, however, we focus specifically on fine-mapping, where performance is particularly sensitive to LD discrepancy.

Effective and ethical sharing of LD information remains a major challenge as direct sharing of individual-level genotype data is restricted due to privacy and ethical concerns Erlich [2014]). While pre-computed LD matrices have been shared in the past, this practice has never become the standard as these are often large, inflexible, and limited to fixed SNP sets defined by the data sharer and estimating LD for an alternative subset is impossible without returning to the original genotypes. As a practical workaround, researchers often rely on external reference panels to estimate LD. However, discrepancies between the true correlation structure and that inferred from such reference datasets can introduce biases, leading to false positive signals, particularly when strong-effect SNPs are present in fine-mapping studies (Price [2006], Schaid [2018]). Even minor mismatches in LD estimation can propagate into downstream analyses, compromising biological interpretations and, ultimately, the conclusions drawn from these studies. To overcome these limitations, we introduce SAFE-LD (Shrinkage and Anonymization Framework for LD Estimation) — a novel method that transforms raw genotype data into a set of compact vectors, stored in genotype dosage format, with one vector representing each SNP. We refer to the per-SNP, dosage-formatted vectors derived from z-score correlations as SAFE-LD surrogate vectors (‘SAFE-LD surrogates’). These are not genotypes; they are LD-preserving proxies intended solely for estimating correlations/LD and for LD-dependent analyses and could not be interpreted as allele counts or used for allele-frequency–based analyses. These surrogate vectors are stored in a vcf or plink pgen format file based on data from a subset of N sampled individuals and which can be used to faithfully recapitulate the true LD structure. By achieving an almost perfect match with the true correlation patterns observed in the original genotype data, SAFE-LD enables downstream analyses, including fine-mapping, to be performed with high accuracy. Importantly, this method safeguards participant privacy while offering a privacy-preserving, flexible, and efficient solution for disseminating LD information, resolving a key bottleneck in modern genetic analysis workflows. Finally we provide a tool to quickly transform any vcf containing dosage information into their corresponding SAFE-LD surrogates.

## 2 Results

### The SAFE-LD approach

The idea behind SAFE-LD is to use the correlation between the linear regression effect estimates coming from the regression of a large number of traits against the same SNPs. A similar approach has been used for an orthogonal problem where GWAS summary stats have been used to estimate the phenotypic correlation between two traits. This approach requires that there is no true correlation between the phenotypes and the SNPs so that the correlation between effect estimates is due solely to the original phenotypic correlation and the complete sample overlap. To overcome this issue, instead of using real data, we generate a large set of randomly generated phenotypes against which the effect size of the regression is estimated. This produces for each SNP a vector of effect size estimates the same size of the number of simulated sets. If this number is big enough, the correlation between the vectors of two different SNPs will be the same as the correlation between the original genotypes for the two SNPs. These vectors are then scaled between 0 and 2 to mimic dosages from imputation and can be stored in any compatible format (ie plink2 or vcf) and used for correlation purposes only, exactly as though they were the genotypes from a population of size equal to the number of randomly simulated traits (SAFE-LD surrogates) and that can be directly plugged in in any existing pipeline or workflow. Given the way they are produced the SAFE-genotypes retain no personal information and even allelic frequencies are lost in the process. However they can still be used for downstream analyses based on summary statistics that require LD information.

### 2.1 Accurate LD matrix reconstruction from z-scores

We first evaluated the ability of SAFE-LD to reconstruct the true internal LD matrix. The accuracy of the reconstructed LD matrix, measured by Pearson correlation with the true LD matrix, depended primarily on the number of simulated phenotypes (*P*) used for the estimation. Our simulations showed that accuracy increases with *P*, with correlations exceeding 0.97 when *P* ≥ 1000 and plateauing around *P* = 3000, where the correlation reached 0.995 (Figure 1a, Supp. Table 2). Conversely, the method’s performance was remarkably stable across different sample sizes (*N*). For a fixed *P* = 3000, the correlation between SAFE-LD and true LD remained high (>0.99) for sample sizes ranging from 200 to 30,000 individuals (Figure 1b, Supp. Table 3). Furthermore, SAFE-LD performed consistently across all 22 autosomes, confirming its invariance to different genomic regions (Figure 1, Supp. Table 5).

**Figure 1.**
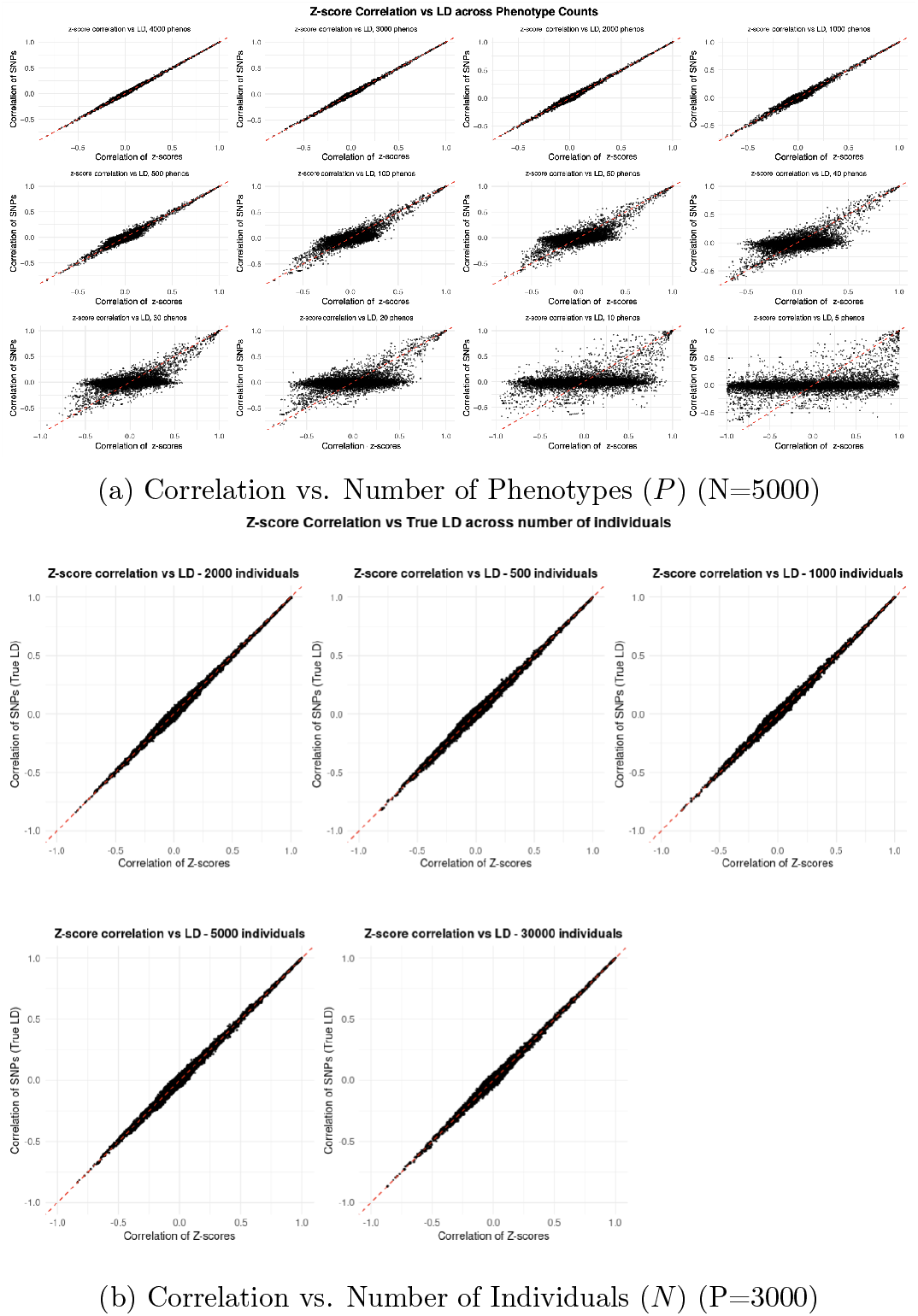
Performance of SAFE-LD in Reconstructing the LD Matrix (A–B). (A) Correlation vs. number of phenotypes (*P*). (B) Correlation vs. number of individuals (*N*).

**Figure 2.**
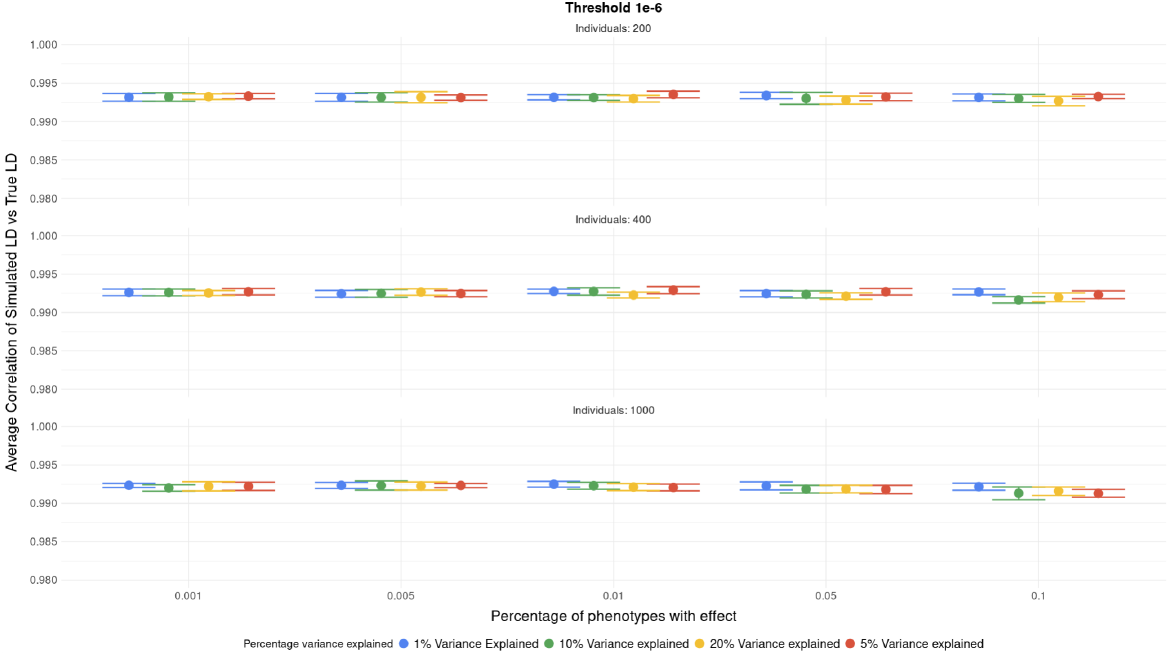
Performance of SAFE-LD in Reconstructing the LD Matrix (C). (C) Correlation under SAFE-LDss across varying *N*, effect sizes (PVE), and proportions of phenotypes with effect [cite: 362].

### 2.2 SAFE-LD enables accurate fine-mapping

Although high correlation indicates a good global approximation of the LD matrix, the ultimate test is its performance in downstream applications. Fine-mapping of strong genetic effects are known to be particularly sensitive to LD discrepancies and thus represent an ideal test for SAFE-LD accuracy. We used the SuSiE fine-mapping framework to compare the performance of SAFE-LD against gold-standard internal LD and to external LD reference panel. In order to compare SAFE-LD to the best-case scenario of cohort-matched external LD where a subset of the cohort is used as reference, we took 30,000 randomly selected white, British individuals present in UKB data and performed a random split, using 2/3 of the data for internal LD and 1/3 of the data for external LD. Performance was evaluated using coverage and power. We first assessed fine-mapping performance using SAFE-LD matrices generated systematically varying the number of individuals (*N*) and phenotypes (*P*) evaluating also the relationship in fine-mapping accuracy and the *N* to P.

Accuracy was evaluated in two ways: power (the ability to detect a true signal) and coverage (whether all detected credible sets contain a true effect). Fig 4 shows the results of the simulations for N equal to 1000, 5000 and 10000 samples. Overall SAFE-LD performed better than the cohort matched LD reference panels both in terms of coverage and power. Relative to internal LD, coverage was comparable (sometimes slightly higher) with a modest reduction in power; power still exceeded the cohort-matched external panel. A different situation presented itself when the number of individuals was greater than or equal to the number of phenotypes (*N ≥ P*), with 10,000 individuals. When (*P* ≪ *N*), SAFE-LD performed poorly, even worse than external LD (Figure 3a, Supp. Tables 6-7) in terms of both coverage and power. However, as *P* approached *N*, performance rapidly improved, in particular there was a turning point at *P* = *N/*2. When *P* ≈ *N*, SAFE-LD achieved coverage and power comparable to, and sometimes exceeding, internal LD (Figure 3, Supp. Tables 6-7) and both coverage and power achieved values > 0.9.

**Figure 3.**
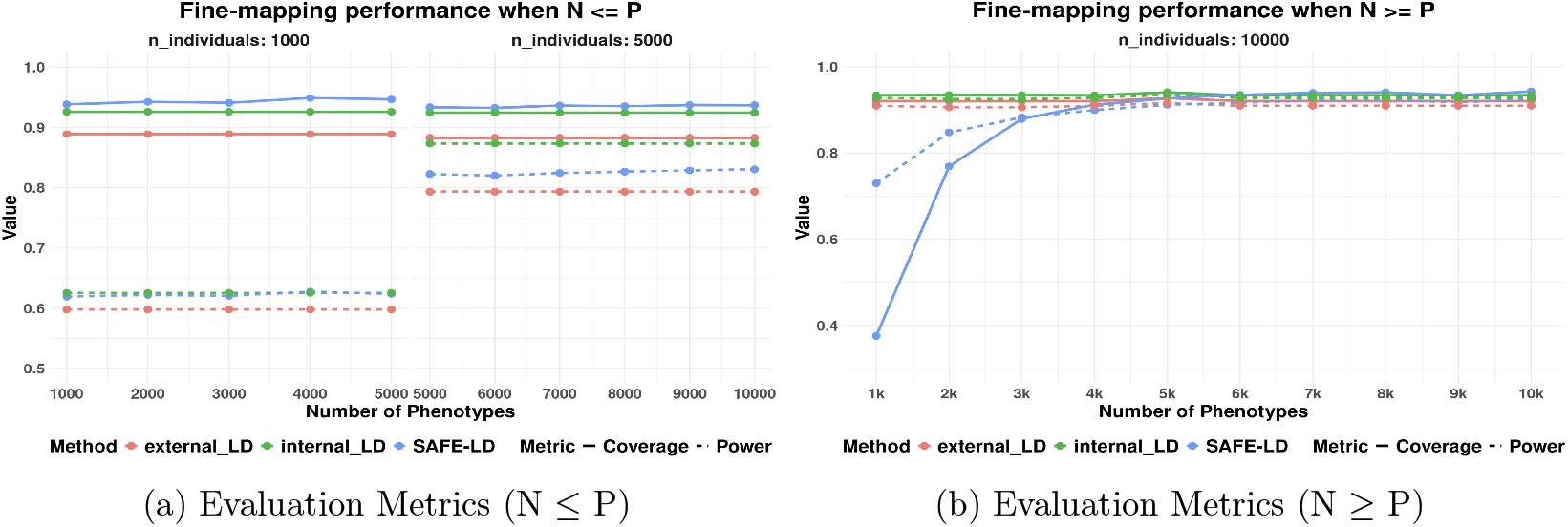
Fine-mapping performance of SAFE-LD vs. Internal and External LD. (a) Mean coverage and mean power when *N ≤ P* . (b) Mean coverage and mean power when *N ≥ P* . Results are averaged over 10 loci with a simulated effect vector of (0.005, 0.05, 0.2). External LD is from a subset of the same cohort as internal LD to mimic ancestry-matched LD.

**Figure 4.**
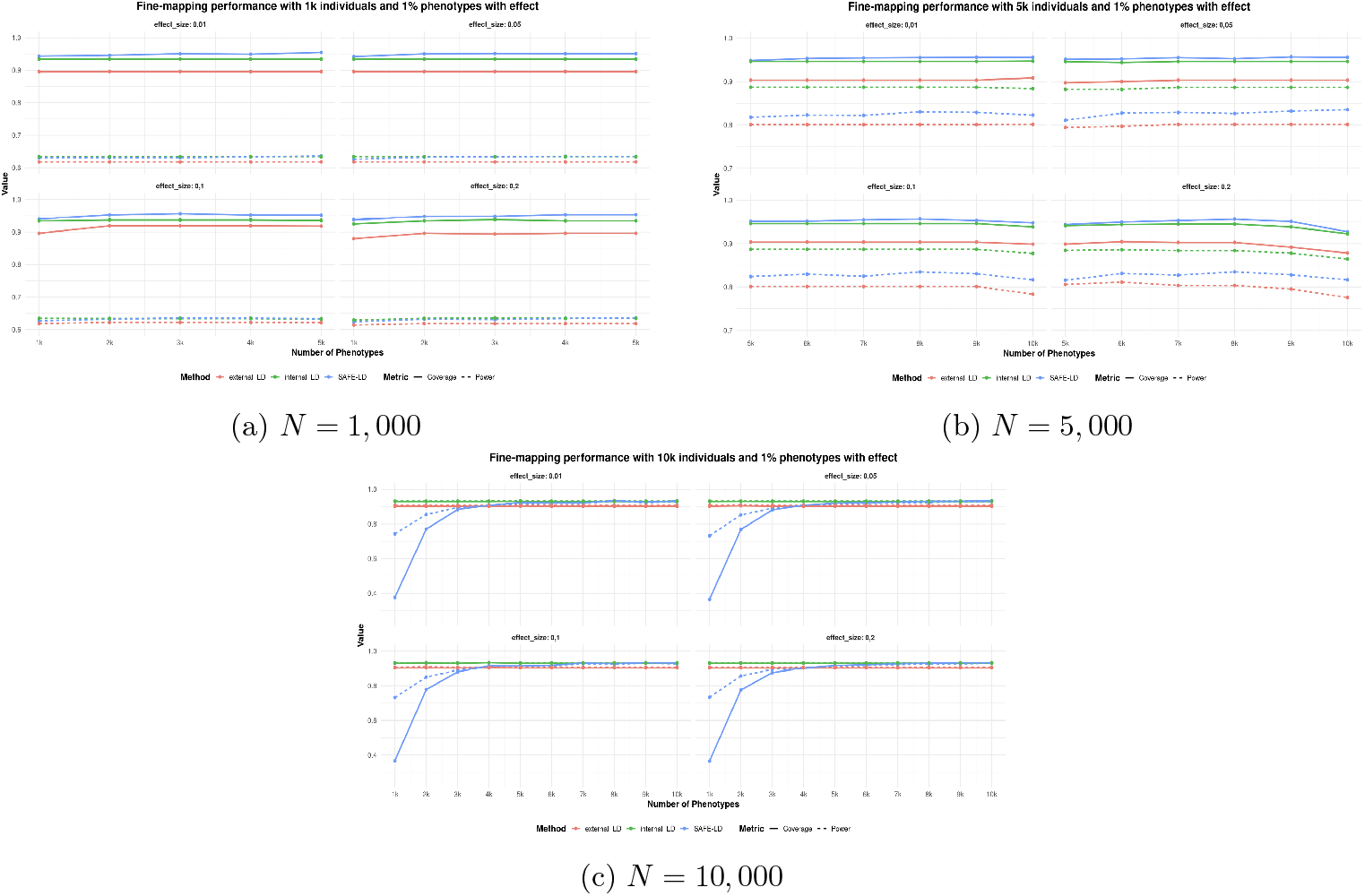
Fine-mapping performance of SAFE-LDss across different N with 1% of phenotypes with effect. Mean coverage and power are plotted as a function of the number of phenotypes (*P*). Results are shown for three scenarios of causal architecture and averaged across four different effect sizes

#### 2.2.1 SAFE-LDss

SAFE-LD relies on estimating the regression coefficient estimates between a number of randomly simulated phenotypes and existing genotypes. Similar information could be obtained from existing summary statistics as long as: a) a large enough number of traits have been analyzed on the same people as in the case of omic datasets b) there is no true association between the examined SNPs and any of the traits. The second condition can be achieved potentially by assuming that traits where no SNP amongst the ones of interest show a p-value smaller than a certain threshold. We thus decided to test which threshold would minimise bias. After testing different possible thresholds, *p <* 1×10^−6^ to filter phenotypes showed good performance while retaining a large number of traits. SAFE-LDss robustly reconstructed the LD matrix even in the presence of sparse causal effects of varying magnitude. The accuracy of the reconstruction was consistently high (Pearson correlation > 0.99) across all tested scenarios, including those with up to 10% of phenotypes having a causal SNP explaining as much as 20% of the variance (Figure 4C). These results confirm that a simple thresholding strategy is effective at removing confounding from true genetic effects, allowing the underlying LD structure to be accurately estimated from z-score correlations. We then evaluated if SAFE-LDss would perform as well as SAFE-LD also in the fine-mapping task. Similarly to SAFE-LD, when *N* = 10, 000 and *P* ≪ *N* (e.g., *P* = 1000), coverage was poor (<0.4), highlighting the need for sufficient phenotypic diversity (Figure 4A, Supp. Table 10). However, performance improved dramatically as *P* increased. When *P ≈ N* (*N* = *P* = 10, 000), SAFE-LDss achieved both coverage and power greater than 0.925 across all tested effect sizes and proportions of causal phenotypes (Figure 4, Supp. Table 10). In this optimal setting, SAFE-LD’s performance was virtually indistinguishable from internal LD and vastly superior to external LD.

At smaller sample sizes (*N* = 1000 and *N* = 5000), SAFE-LDss consistently provided coverage as good as or better than internal LD (Supp. Tables 8-9). Power was more sensitive to sample size, with *N* = 1000 being insufficient to achieve high power regardless of the LD matrix used (max power ∼0.64) (Supp. Table 8). At *N* = 5000, power improved substantially (to >0.8), and SAFE-LD consistently outperformed external LD, though it did not match the power of internal LD (Supp. Table 9).

These comprehensive tests show that when a sufficient number of phenotypes are available (*P ≈ N*) and sample size is around 10,000, SAFE-LDss robustly estimates LD from summary statistics and enables fine-mapping with an accuracy that rivals the use of gold-standard internal LD. At sample sizes of around 5,000, SAFE-LDss consistently outperforms external LD and is a superior alternative, but it does not match the performance of internal LD. At sample sizes of around 1,000, all three LD matrices achieve low power, so we can conclude that this is due to the sample size rather than the methods itself.

## 3 Discussion

In this study, we introduced SAFE-LD, a novel framework for estimating LD matrices directly from summary statistics, thereby circumventing the need for individual-level genotype data. Our results show that SAFE-LD can accurately reconstruct internal LD matrices and, when applied to fine-mapping, achieves performance comparable to the gold standard of using internal LD and substantially outperforms the common practice of using external LD panels even under the best possible condition where the reference dataset is a subset of the original cohort.

The core strength of SAFE-LD lies in its ability to leverage the correlation structure of beta estimates across numerous phenotypes. Unsurprisingly the accuracy of LD reconstruction is highly dependent on the number of phenotypes (*P*) used, with performance plateauing when *P* is in the thousands. This is likely due to the fact that exploiting the correlation between the betas is extremely noisy and thus needs large numbers, however the reliance on simulated phenotypes effectively removes this limitation. Our fine-mapping analyses revealed that the optimal performance is achieved when the number of phenotypes is close to the number of individuals in the study (*P ≈ N*). While increasing *N* consistently improves fine-mapping power and coverage, increasing *P* beyond *N* offers diminishing returns. This provides a practical guideline for researchers: to reliably substitute internal LD, one should ideally use summary statistics from a number of traits comparable to the GWAS sample size. LD-correlation appears to plateau once P reaches the low thousands (≈ 0.995 by *P ≈* 3000), but the P/N requirement for optimal fine-mapping may differ at larger N (e.g., *N ≈* 500k) and merits empirical study. A reasonably elevated sample size is also needed in order to achieve both high coverage and high power. High coverage can be established with as few as 1,000 individuals, but high power needs a sample size of more than 5,000 individuals under the parameters selected for simulations, however the power is close to internal LD. Despite this slight loss in power SAFE-LD still outperforms external LD and thus represents the best alternative to having direct access to internal genotypes. When we consider coverage SAFE-LD slightly outperforms even internal LD compensating the loss in power with a reduction in false discoveries.

**Table 1:** Summary of results demonstrating consistency across two methods.

| Sample Size ($N$) | Coverage | Power | Conclusion |
| --- | --- | --- | --- |
| 1,000 | ✓ | ● | $N$ too small, insufficient for high power |
| 5,000 | ✓ | ● | Power improves with $N$, but still lower than desired and Internal still best |
| 10,000($P < N/2$) | ✗ | ✗ | For $P \ll N$, both power and coverage struggle. |
| 10,000($P \geq N/2$) | ✓ | ✓ | As $P$ approaches $N$, high coverage and power achieved. SAFE-LD matches gold-standard Internal and outperforms External. |

The presented results should be evaluated also considering the fact we have benchmarked our approach against the best possible external reference dataset (a subset of the same cohort). In practice, reference panels are often less well matched to the target cohort, which inflates false credible sets and yields less reliable results. SAFE-LD and SAFE-LDss show broadly similar patterns; in our experiments, the settings where SAFE-LD outperformed the external panel and approached internal LD were similar for both, though not identical in magnitude. This is positive as it highlights that our method is applicable in both the scenario where researchers have individual level genotypic data (allowing them to share LD much more easily) and also in the case where researches only have GWAS (allowing them to capture LD easily and accurately). This result also shows that the method is robust to genetic effects as strong as 0.2 present in up to 10% of SNPs, meaning that it can be applied to most GWAS with no problem.

Our work also highlights how important having precise LD is for finemapping purposes While a high correlation (>0.99) was achieved with *P ≥* 3000, this did not always translate to optimal fine-mapping performance. When the two were compared directly, most discrepancies arose for weak correlations near zero, suggesting that even slight deviation from the true values may have great impact on downstream analyses. This is important as when LD matrices are shared they are generally sparsified and values below a certain threshold are excluded as considered unimportant. Our results show that this is not the case and that it is important to retain the ability to retaining (or reconstructing) the full dense LD. The study has some limitations that open avenues for future research. First, our model assumes that the phenotypes used for LD estimation in SAFE-LDss are uncorrelated. In practice, many traits are phenotypically correlated, which could potentially introduce bias into the LD estimates. Investigating the impact of such correlations and developing methods to correct for them is an important next step. Second, our SAFE-LDss employs a hard p-value threshold to exclude phenotypes with genetic effects. This approach, while effective, could be replaced by more sophisticated regularization or weighting techniques that down-weight, rather than discard, signals from associated SNPs, potentially leading to a more stable and robust estimation.

In conclusion, SAFE-LD provides a powerful and practical solution to a major bottleneck in genetic analysis. By eliminating the dependence on inaccessible individual-level data, it empowers researchers to perform high-accuracy fine-mapping and other LD-dependent analyses. It allows researcher to share LD easily and without any privacy concern for the cohort participants. Furthermore, SAFE-LD offers a significant logistical advantage by obviating the need to store, manage, and share massive LD matrix files as it is compatible with most existing softwares. As the number of available GWAS summary statistics continues to grow, methods like SAFE-LD will become increasingly vital tools for unlocking new insights into the genetic architecture of complex traits.

## 4 Methods

### 4.1 Statistical Foundation

The SAFE-LD method is based on the principle that, under the null hypothesis of no genetic effect, the correlation between GWAS z-scores for any two SNPs is an unbiased estimator of the linkage disequilibrium between them. For a given trait *t*, the z-score for SNP *j* in a study of *N* individuals is 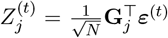, where **G**_*j*_ is the standardized genotype vector for SNP *j* and ***ε***^(*t*)^ is the vector of standardized residuals (which, under the null, is the phenotype vector **y**^(*t*)^). The expected product of z-scores for two SNPs, *i* and *j*, is:

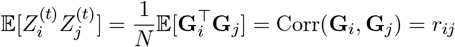

where *r*_*ij*_ is the Pearson correlation coefficient (LD) between the genotypes of SNP *i* and SNP *j*[cite: 163]. In practice, we estimate *r*_*ij*_ by computing the empirical Pearson correlation of the z-score vectors for SNPs *i* and *j* across a large number of traits (*T*):

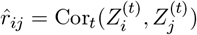

The precision of this estimator increases with the number of traits used.

### 4.2 SAFE-LD Methods

We developed two methods based on this principle:

1. **SAFE-LD:** This method simulates an ideal scenario where a user has access to individual-level genotype data but wishes to create a privacy-preserving LD matrix. A large number of null phenotypes are generated (e.g., from a standard normal distribution, *Y ∼* 𝒩(0, 1)) and regressed against the genotypes one SNP at a time to generate GWAS summary statistics. The resulting z-scores (calculated as 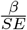) are then correlated to compute the z-score correlation matrix, which serves as the SAFE-LD matrix.
2. **SAFE-LDss:** This method is designed for the common scenario where only pre-existing GWAS summary statistics for multiple traits are available. Since these traits may have true genetic associations, the z-score correlation would be biased by the effects. To mitigate this, we apply a p-value thresholding strategy. For each trait, we test for genetic effects across all SNPs in the region of interest. If any SNP shows a significant association (e.g., *p <* 1×10^−6^), the entire trait is excluded from the analysis. The SAFE-LD matrix is then computed by correlating the z-scores from the remaining “null” traits.

### 4.3 Simulation and Data

All simulations and analyses were conducted using genotype data from UK Biobank (Biobank. Our primary cohort consisted of 30,000 unrelated, white British individuals. Genotypes were filtered for minor allele frequency (MAF > 0.01) and Hardy-Weinberg equilibrium. For external LD comparison, we used either a randomly-selected one-third of this cohort (10,000 individuals) to simulate a best-case, ancestry-matched external panel.

Phenotypes were simulated under two conditions:

1. **Null simulations**, where phenotypes were drawn from 𝒩 (0, 1)
2. **Causal effect simulations**, where for a subset of phenotypes, a causal SNP was chosen to explain a certain proportion of variance (PVE)

GWAS summary statistics were generated using PLINK2’s (Shaun Purcell, Christopher C Chang [2015]) ‘–glm’ function.

### 4.4 Fine-Mapping and Evaluation

We used the SuSiE (Sum of Single Effects) (Gao Wang. Abhishek Sarkar [2020]) regression framework, implemented in the ‘susieR’ R package, to perform fine-mapping [Wang et al., 2020, Gao Wang. Abhishek Sarkar, 2020]. We simulated causal effects for 10 distinct genomic loci Supp. Table 1) with a complex effect vector of (PVE = 0.005, 0.05, 0.2) to represent a range of weak to strong signals.

Fine-mapping performance was evaluated for four LD matrix types:

1. **Internal LD:** The gold standard, computed directly from the genotype data of the main study cohort.
2. **External LD:** Computed from a separate, non-overlapping reference panel.
3. **SAFE-LD:** Generated from simulated null phenotypes.
4. **SAFE-LDss:** Generated from phenotypes with sparse causal effects using a p-value threshold.

Performance was measured by **coverage** (the proportion of 95% credible sets that contain a true causal SNP) and **power** (the proportion of true causal SNPs that are included in any credible set).

## 5 Data and Code Availability

All code used for this project as well as R script for SAFE-LDss is publicly available on GitHub: https://github.com/gidiesse/SAFE-LD. A high-performance C++ implementation of the SAFE-LD (Linkage Disequilibrium) simulator for generating synthetic genomic data from VCF files is available at: https://github.com/davidebolo1993/safeld.

## 6 Acknowledgements

This research was conducted using the UK Biobank Resource (Biobank) under Application Number 102297. This work was made possible through the support and hosting of Human Technopole.

## Supplementary Information

## Supplementary Figures

**Figure 5.**
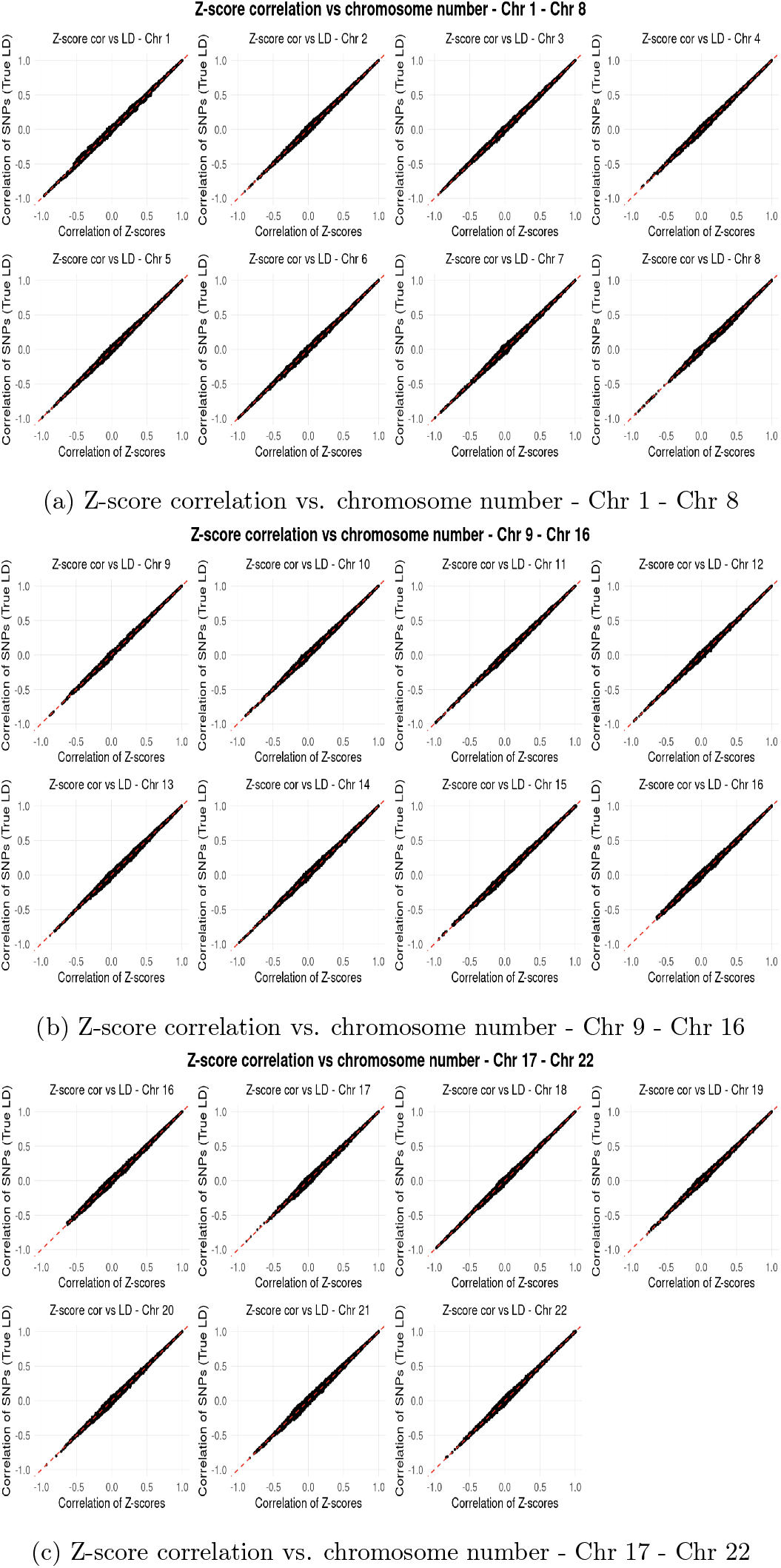
Invariance of SAFE-LD to Genomic Region. Scatter plots comparing true LD (x-axis) and SAFE-LD (y-axis). In all cases, the Pearson correlation is >0.99, demonstrating consistent performance across the genome. Simulations were run with N=30,000 and P=3000. (a) Chromosomes 1 - 8 (b) Chromosomes 9 - 16 (c) Chromosomes 17 - 22

**Figure 6.**
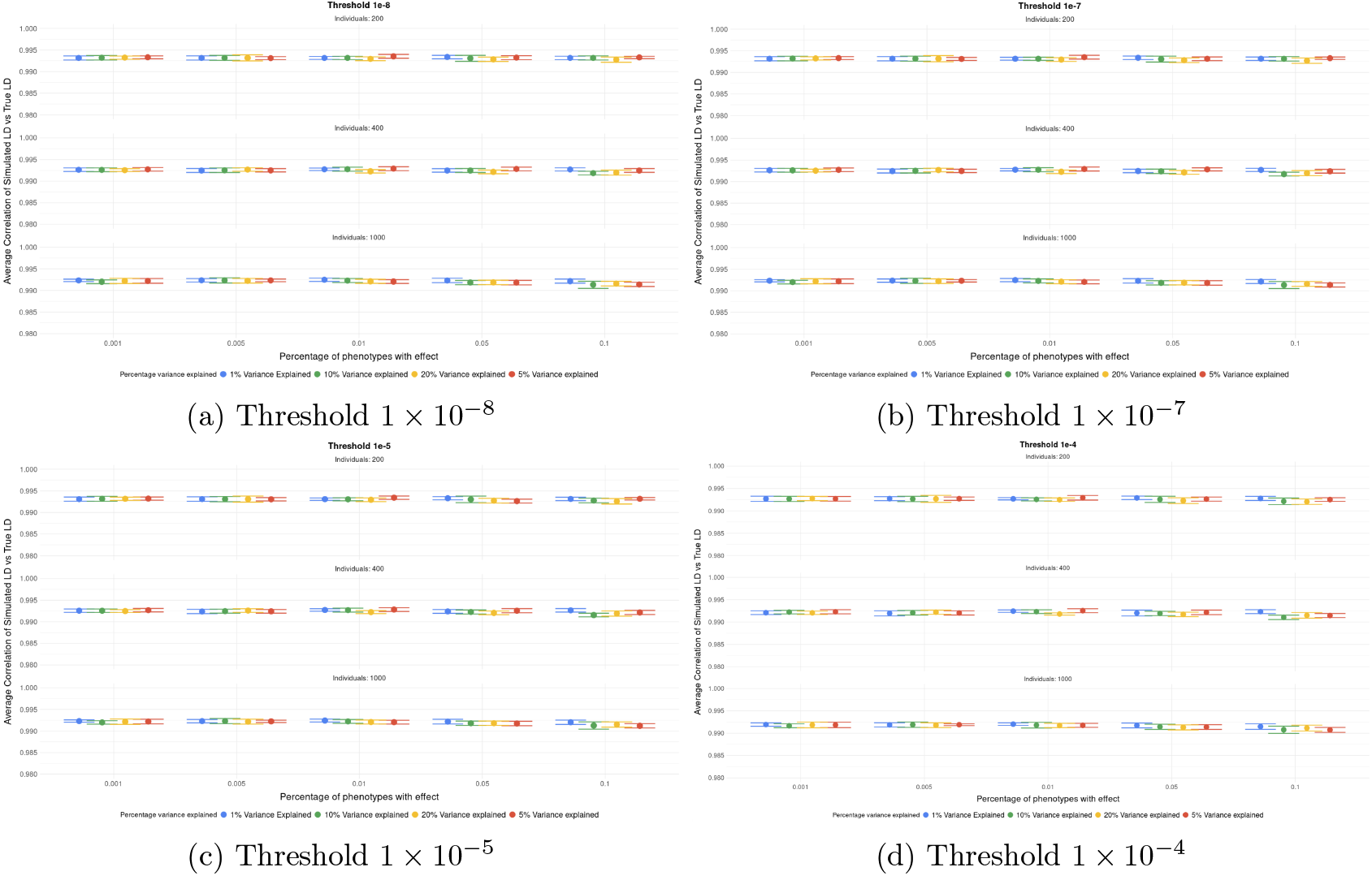
Performance of SAFE-LDss with Different P-value Thresholds. Pearson correlation between true LD and SAFE-LDss at four different p-value thresholds: (a) 1 *×* 10^−8^, (b) 1 *×* 10^−7^, (c) 1 *×* 10^−5^, and (d) 1 *×* 10^−4^ Performance is robust across all tested thresholds.

**Figure 7.**
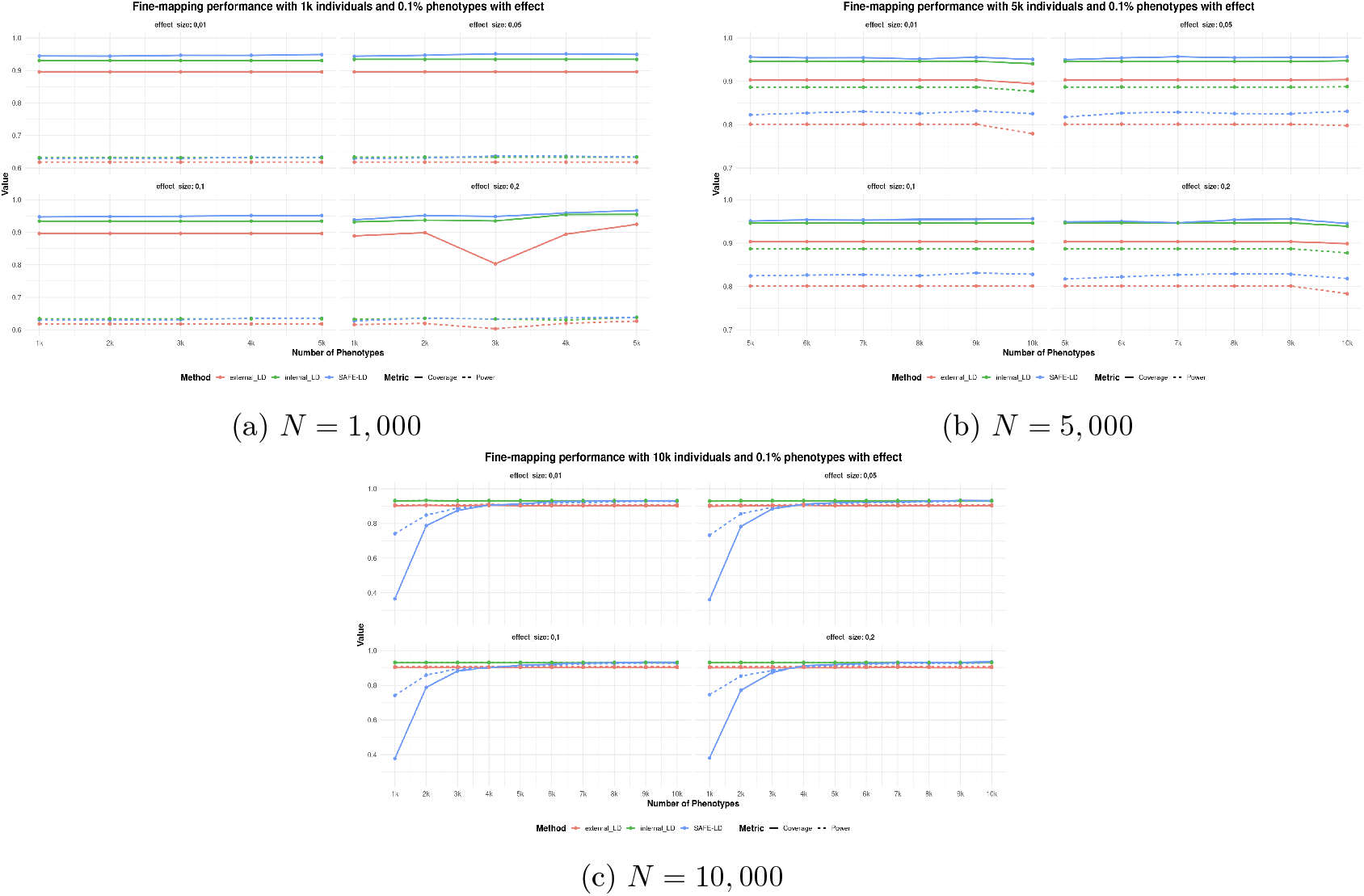
Fine-mapping performance of SAFE-LD across different N with 0.1% of phenotypes with effect. Mean coverage and power are plotted as a function of the number of phenotypes (*P*). Results are shown for three scenarios of causal architecture and averaged across four different effect sizes. (a) Fine-mapping on 1,000 individuals. (b) Fine-mapping on 5,000 individuals. (c) Fine-mapping on 10,000 individuals.

**Figure 8.**
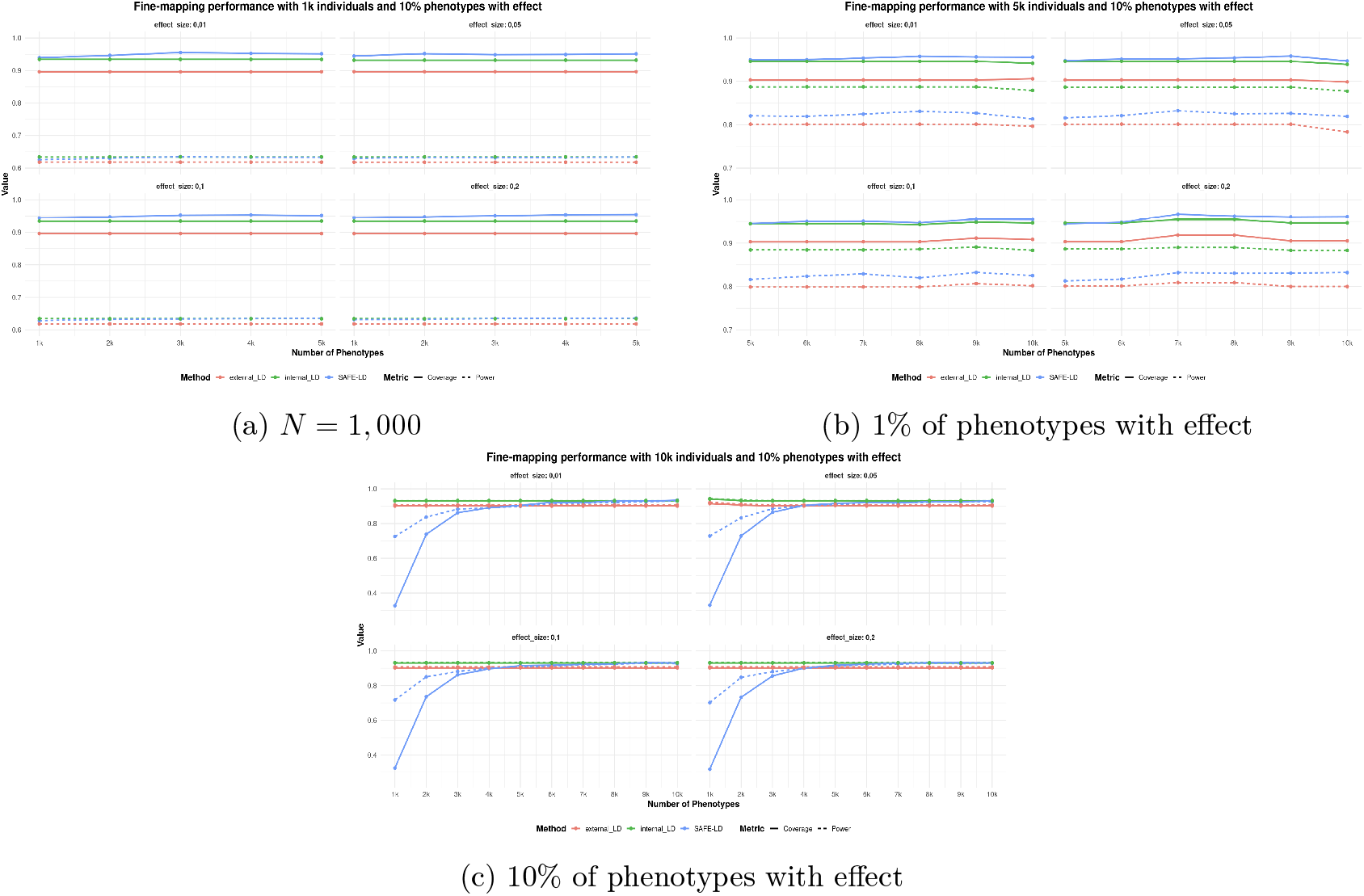
Fine-mapping performance of SAFE-LD across different N with 10% of phenotypes with effect. Mean coverage and power are plotted as a function of the number of phenotypes (*P*). Results are shown for three scenarios of causal architecture and averaged across four different effect sizes. (a) Fine-mapping on 1,000 individuals. (b) Fine-mapping on 5,000 individuals. (c) Fine-mapping on 10,000 individuals.

## Supplementary Tables

**Supplementary Table 1.** Loci used for fine-mapping simulations.

| Locus | Chr | Start | End | Locus | Chr | Start | End | Locus | Chr | Start | End |
| --- | --- | --- | --- | --- | --- | --- | --- | --- | --- | --- | --- |
| Locus 1 | 6 | 134597 | 135597 | Locus 34 | 3 | 23793 | 24793 | Locus 68 | 7 | 49860 | 50860 |
| Locus 2 | 19 | 44745 | 45745 | Locus 35 | 4 | 54042 | 55042 | Locus 69 | 8 | 41183 | 42183 |
| Locus 3 | 6 | 108741 | 109745 | Locus 36 | 5 | 787 | 1787 | Locus 70 | 15 | 77744 | 78744 |
| Locus 4 | 2 | 23175 | 24175 | Locus 37 | 6 | 41457 | 42457 | Locus 71 | 16 | -318 | 681 |
| Locus 5 | 2 | 27008 | 28008 | Locus 38 | 8 | 41273 | 42273 | Locus 72 | 17 | 43752 | 44752 |
| Locus 6 | 12 | 110947 | 111947 | Locus 39 | 9 | 21643 | 22643 | Locus 73 | 17 | 77629 | 78629 |
| Locus 7 | 1 | 158061 | 159061 | Locus 40 | 10 | 24412 | 25412 | Locus 74 | 19 | 3985 | 4985 |
| Locus 8 | 20 | 1441 | 2441 | Locus 41 | 10 | 48667 | 49667 | Locus 75 | 19 | 16642 | 17642 |
| Locus 9 | 9 | 132762 | 133762 | Locus 42 | 10 | 68835 | 69835 | Locus 76 | 1 | 3275 | 4275 |
| Locus 10 | 1 | 198532 | 199532 | Locus 43 | 11 | 100080 | 101080 | Locus 77 | 2 | 8116 | 9116 |
| Locus 11 | 6 | 134456 | 135456 | Locus 44 | 11 | 107654 | 108654 | Locus 78 | 2 | 23390 | 24390 |
| Locus 12 | 8 | 124968 | 125968 | Locus 45 | 11 | 116252 | 117252 | Locus 79 | 2 | 112583 | 113583 |
| Locus 13 | 9 | 88278 | 89278 | Locus 46 | 12 | 110948 | 111948 | Locus 80 | 3 | 122850 | 123850 |
| Locus 14 | 17 | 44912 | 45912 | Locus 47 | 14 | 24476 | 25476 | Locus 81 | 3 | 141973 | 142973 |
| Locus 15 | 1 | 117097 | 118097 | Locus 48 | 16 | 88271 | 89271 | Locus 82 | 3 | 196275 | 197275 |
| Locus 16 | 2 | 42725 | 43725 | Locus 49 | 17 | 16449 | 17449 | Locus 83 | 6 | 25604 | 26604 |
| Locus 17 | 5 | 605 | 1605 | Locus 50 | 1 | 45319 | 46319 | Locus 84 | 6 | 41439 | 42439 |
| Locus 18 | 9 | 132487 | 133487 | Locus 51 | 2 | 210176 | 211176 | Locus 85 | 6 | 139007 | 140007 |
| Locus 19 | 9 | 132489 | 133489 | Locus 52 | 6 | 15748 | 16748 | Locus 86 | 7 | 92108 | 93108 |
| Locus 20 | 11 | 61303 | 62303 | Locus 53 | 6 | 25593 | 26593 | Locus 87 | 8 | 29922 | 30922 |
| Locus 21 | 12 | 3721 | 4721 | Locus 54 | 10 | 62845 | 63845 | Locus 88 | 9 | 132753 | 133753 |
| Locus 22 | 18 | 62754 | 63754 | Locus 55 | 14 | 40079 | 41079 | Locus 89 | 9 | 133530 | 134530 |
| Locus 23 | 20 | 8124 | 9124 | Locus 56 | 14 | 64513 | 65513 | Locus 90 | 10 | 44393 | 45393 |
| Locus 24 | 20 | 32038 | 33038 | Locus 57 | 15 | 64929 | 65929 | Locus 91 | 10 | 44957 | 45957 |
| Locus 25 | 22 | 21062 | 22062 | Locus 58 | 19 | 12370 | 13370 | Locus 92 | 10 | 99012 | 100012 |
| Locus 26 | 1 | 204671 | 205671 | Locus 59 | 20 | 3662 | 4662 | Locus 93 | 11 | 113605 | 114605 |
| Locus 27 | 1 | 247376 | 248376 | Locus 60 | 1 | 203181 | 204181 | Locus 94 | 16 | 29592 | 30592 |
| Locus 28 | 18 | 45776 | 46776 | Locus 61 | 3 | 128083 | 129083 | Locus 95 | 18 | 43894 | 44894 |
| Locus 29 | 22 | 36567 | 37567 | Locus 62 | 4 | 109415 | 110415 | Locus 96 | 19 | 2680 | 3680 |
| Locus 30 | 22 | 50025 | 51025 | Locus 63 | 4 | 143550 | 144550 | Locus 97 | 19 | 16641 | 17641 |
| Locus 31 | 1 | 23014 | 24014 | Locus 64 | 5 | 782 | 1782 | Locus 98 | 19 | 32764 | 33764 |
| Locus 32 | 2 | 240071 | 241071 | Locus 65 | 6 | 43289 | 44289 | Locus 99 | 20 | 56915 | 57915 |
| Locus 33 | 3 | 11694 | 12694 | Locus 66 | 6 | 89726 | 90726 | Locus 100 | 22 | 43429 | 44429 |
|  |  |  |  | Locus 67 | 6 | 139011 | 140011 |  |  |  |  |

**Supplementary Table 2.** Correlation vs. Number of Phenotypes (SAFE-LD).

| No. of Phenotypes | Correlation |
| --- | --- |
| 4000 | 0.99607 |
| 3000 | 0.99479 |
| 2000 | 0.99272 |
| 1000 | 0.98393 |
| 500 | 0.96726 |
| 100 | 0.86864 |

**Supplementary Table 3.**
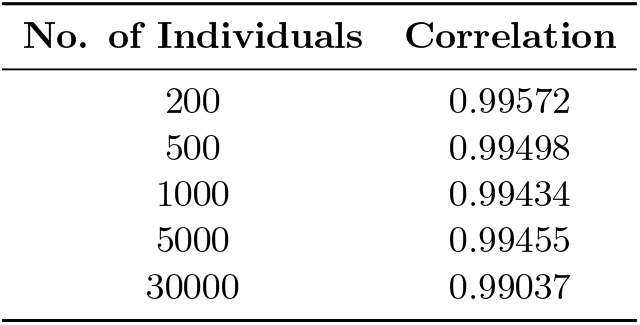
Correlation vs. Number of Individuals (SAFE-LD).

| No. of Individuals | Correlation |
| --- | --- |
| 200 | 0.99572 |
| 500 | 0.99498 |
| 1000 | 0.99434 |
| 5000 | 0.99455 |
| 30000 | 0.99037 |

**Supplementary Table 4.** Genomic loci used for fine-mapping simulations.

| Locus | Chromosome | Start | End |
| --- | --- | --- | --- |
| Locus 1 | 6 | 134597 | 135597 |
| Locus 2 | 19 | 44745 | 45745 |
| Locus 3 | 6 | 108741 | 109745 |
| Locus 4 | 2 | 23175 | 24175 |
| Locus 5 | 2 | 27008 | 28008 |
| Locus 6 | 12 | 110947 | 111947 |
| Locus 7 | 1 | 158061 | 159061 |
| Locus 8 | 20 | 1441 | 2441 |
| Locus 9 | 9 | 132762 | 133762 |
| Locus 10 | 1 | 198532 | 199532 |

**Supplementary Table 5.** Correlation between real LD and simulated LD for a single iteration.

| Chromosome | Correlation | Chromosome | Correlation |
| --- | --- | --- | --- |
| 1 | 0.9984714 | 12 | 0.9977469 |
| 2 | 0.9957691 | 13 | 0.9971851 |
| 3 | 0.9989885 | 14 | 0.9979875 |
| 4 | 0.9971892 | 15 | 0.9988324 |
| 5 | 0.9980794 | 16 | 0.9982558 |
| 6 | 0.9992581 | 17 | 0.9933221 |
| 7 | 0.9951583 | 18 | 0.9976377 |
| 8 | 0.9985925 | 19 | 0.9980026 |
| 9 | 0.9991324 | 20 | 0.9948137 |
| 10 | 0.9984078 | 21 | 0.9972096 |
| 11 | 0.9987795 | 22 | 0.9948448 |

**Supplementary Table 6.**
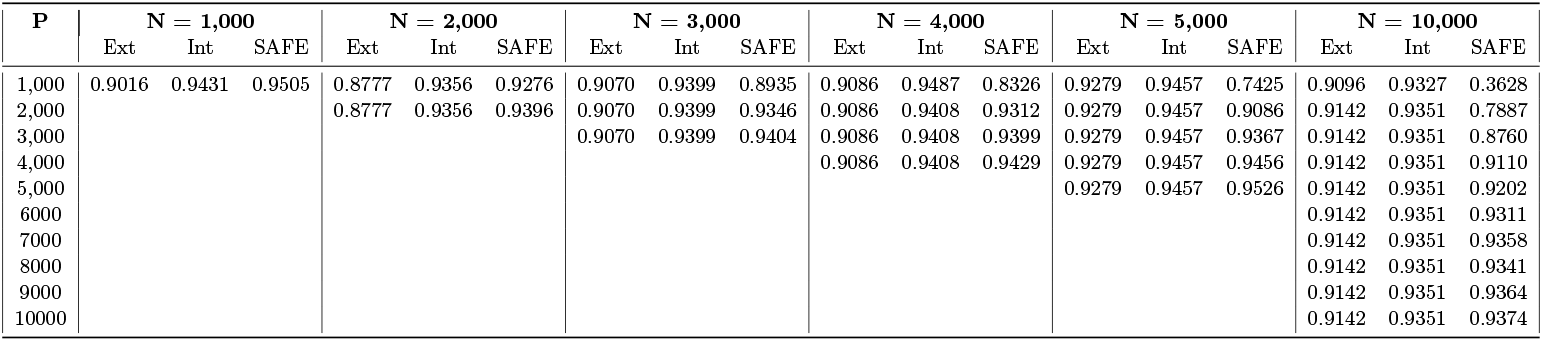
Mean coverage by number of phenotypes and individuals.

**Supplementary Table 7.** Mean power by number of phenotypes and individuals.

| P | N = 1,000 |  |  | N = 2,000 |  |  | N = 3,000 |  |  | N = 4,000 |  |  | N = 5,000 |  |  | N = 10,000 |  |  |
| --- | --- | --- | --- | --- | --- | --- | --- | --- | --- | --- | --- | --- | --- | --- | --- | --- | --- | --- |
|  | Ext | Int | SAFE | Ext | Int | SAFE | Ext | Int | SAFE | Ext | Int | SAFE | Ext | Int | SAFE | Ext | Int | SAFE |
| 1,000 | 0.6144 | 0.6350 | 0.6257 | 0.6394 | 0.6827 | 0.6540 | 0.7033 | 0.7600 | 0.6953 | 0.7514 | 0.8407 | 0.7293 | 0.8117 | 0.8950 | 0.7573 | 0.9104 | 0.9322 | 0.7274 |
| 2,000 |  |  |  | 0.6394 | 0.6827 | 0.6523 | 0.7033 | 0.7600 | 0.7060 | 0.7514 | 0.8350 | 0.7540 | 0.8117 | 0.8950 | 0.7943 | 0.9130 | 0.9333 | 0.8527 |
| 3,000 |  |  |  |  |  |  | 0.7033 | 0.7600 | 0.7017 | 0.7514 | 0.8350 | 0.7613 | 0.8117 | 0.8950 | 0.8160 | 0.9130 | 0.9333 | 0.8900 |
| 4,000 |  |  |  |  |  |  |  |  |  | 0.7514 | 0.8350 | 0.7637 | 0.8117 | 0.8950 | 0.8260 | 0.9130 | 0.9333 | 0.9087 |
| 5,000 |  |  |  |  |  |  |  |  |  |  |  |  | 0.8117 | 0.8950 | 0.8290 | 0.9130 | 0.9333 | 0.9130 |
| 6000 |  |  |  |  |  |  |  |  |  |  |  |  |  |  |  | 0.9130 | 0.9333 | 0.9237 |
| 7000 |  |  |  |  |  |  |  |  |  |  |  |  |  |  |  | 0.9130 | 0.9333 | 0.9240 |
| 8000 |  |  |  |  |  |  |  |  |  |  |  |  |  |  |  | 0.9130 | 0.9333 | 0.9260 |
| 9000 |  |  |  |  |  |  |  |  |  |  |  |  |  |  |  | 0.9130 | 0.9333 | 0.9310 |
| 10000 |  |  |  |  |  |  |  |  |  |  |  |  |  |  |  | 0.9130 | 0.9333 | 0.9307 |

**Supplementary Table 8.**
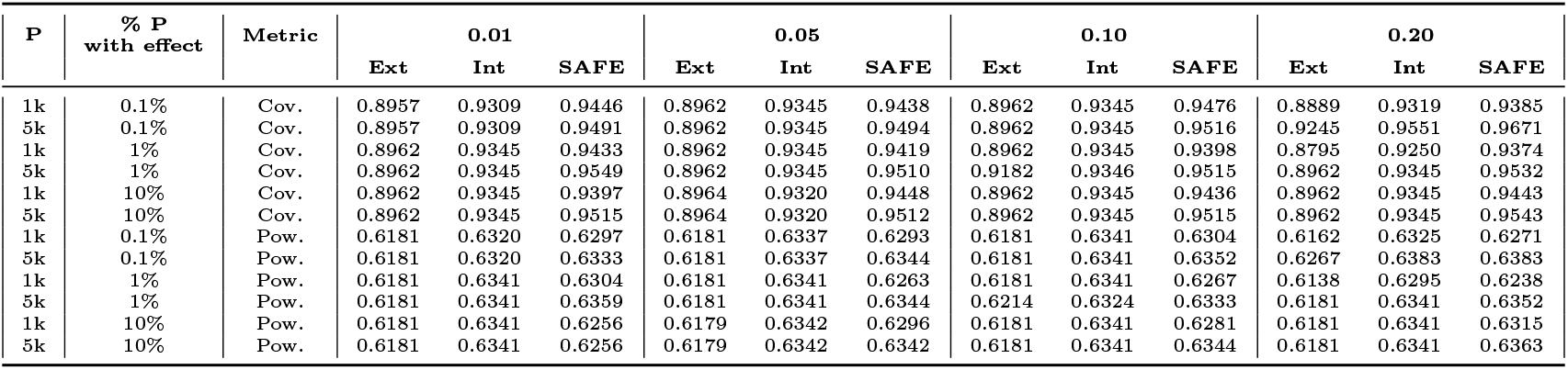
Mean coverage and power for 1,000 and 5,000 phenotypes with 1,000 individuals across four effect sizes.

**Supplementary Table 9.** Mean coverage and power for 5,000 and 10,000 phenotypes with 5000 individuals across four effect sizes.

| P | % P<br>with effect | Metric | 0.01 |  |  | 0.05 |  |  | 0.10 |  |  | 0.20 |  |  |
| --- | --- | --- | --- | --- | --- | --- | --- | --- | --- | --- | --- | --- | --- | --- |
|  |  |  | Ext | Int | SAFE | Ext | Int | SAFE | Ext | Int | SAFE | Ext | Int | SAFE |
| 5k | 0.1% | Cov. | 0.9034 | 0.9465 | 0.9565 | 0.9034 | 0.9459 | 0.9502 | 0.9034 | 0.9462 | 0.9510 | 0.9034 | 0.9466 | 0.9489 |
| 10k | 0.1% | Cov. | 0.8946 | 0.9406 | 0.9509 | 0.9046 | 0.9474 | 0.9565 | 0.9034 | 0.9462 | 0.9564 | 0.8988 | 0.9392 | 0.9453 |
| 5k | 1% | Cov. | 0.9034 | 0.9466 | 0.9488 | 0.8975 | 0.9463 | 0.9517 | 0.9034 | 0.9462 | 0.9514 | 0.8990 | 0.9413 | 0.9441 |
| 10k | 1% | Cov. | 0.9086 | 0.9474 | 0.9562 | 0.9035 | 0.9465 | 0.9562 | 0.8988 | 0.9384 | 0.9478 | 0.8782 | 0.9225 | 0.9273 |
| 5k | 10% | Cov. | 0.9034 | 0.9462 | 0.9498 | 0.9034 | 0.9462 | 0.9479 | 0.9031 | 0.9449 | 0.9451 | 0.9034 | 0.9465 | 0.9446 |
| 10k | 10% | Cov. | 0.9061 | 0.9418 | 0.9560 | 0.8988 | 0.9392 | 0.9471 | 0.9086 | 0.9468 | 0.9548 | 0.9052 | 0.9465 | 0.9608 |
| 5k | 0.1% | Pow. | 0.8011 | 0.8867 | 0.8227 | 0.8011 | 0.8870 | 0.8177 | 0.8011 | 0.8870 | 0.8243 | 0.8011 | 0.8870 | 0.8173 |
| 10k | 0.1% | Pow. | 0.7793 | 0.8773 | 0.8253 | 0.7979 | 0.8878 | 0.8311 | 0.8011 | 0.8870 | 0.8280 | 0.7833 | 0.8775 | 0.8183 |
| 5k | 1% | Pow. | 0.8007 | 0.8870 | 0.8180 | 0.7944 | 0.8824 | 0.8110 | 0.8011 | 0.8867 | 0.8240 | 0.8057 | 0.8842 | 0.8158 |
| 10k | 1% | Pow. | 0.8017 | 0.8838 | 0.8229 | 0.8015 | 0.8867 | 0.8353 | 0.7833 | 0.8775 | 0.8167 | 0.7756 | 0.8644 | 0.8167 |
| 5k | 10% | Pow. | 0.8011 | 0.8873 | 0.8203 | 0.8011 | 0.8867 | 0.8157 | 0.7989 | 0.8843 | 0.8162 | 0.8011 | 0.8867 | 0.8127 |
| 10k | 10% | Pow. | 0.7967 | 0.8792 | 0.8133 | 0.8011 | 0.8775 | 0.8192 | 0.8017 | 0.8833 | 0.8252 | 0.8000 | 0.8833 | 0.8321 |

**Supplementary Table 10.** Mean coverage and power for 1,000, 5,000 and 10,000 phenotypes with 10,000 individuals across four effect sizes.

| P | % P<br>with effect | Metric | 0.01 |  |  | 0.05 |  |  | 0.10 |  |  | 0.20 |  |  |
| --- | --- | --- | --- | --- | --- | --- | --- | --- | --- | --- | --- | --- | --- | --- |
|  |  |  | Ext | Int | SAFE | Ext | Int | SAFE | Ext | Int | SAFE | Ext | Int | SAFE |
| 1k | 0.1% | Cov. | 0.9020 | 0.9301 | 0.3656 | 0.9000 | 0.9286 | 0.3606 | 0.9023 | 0.9301 | 0.3776 | 0.9018 | 0.9301 | 0.3807 |
| 5k | 0.1% | Cov. | 0.9020 | 0.9301 | 0.9128 | 0.9018 | 0.9301 | 0.9204 | 0.9023 | 0.9301 | 0.9155 | 0.9018 | 0.9301 | 0.9202 |
| 10k | 0.1% | Cov. | 0.9020 | 0.9301 | 0.9321 | 0.9018 | 0.9301 | 0.9332 | 0.9023 | 0.9301 | 0.9329 | 0.9018 | 0.9301 | 0.9356 |
| 1k | 1% | Cov. | 0.9020 | 0.9301 | 0.3765 | 0.9020 | 0.9301 | 0.3653 | 0.9023 | 0.9301 | 0.3675 | 0.9020 | 0.9301 | 0.3656 |
| 5k | 1% | Cov. | 0.9031 | 0.9312 | 0.9243 | 0.9020 | 0.9301 | 0.9207 | 0.9023 | 0.9301 | 0.9146 | 0.9020 | 0.9301 | 0.9171 |
| 10k | 1% | Cov. | 0.9020 | 0.9301 | 0.9339 | 0.9020 | 0.9301 | 0.9350 | 0.9023 | 0.9301 | 0.9319 | 0.9020 | 0.9301 | 0.9340 |
| 1k | 10% | Cov. | 0.9018 | 0.9301 | 0.3272 | 0.9152 | 0.9409 | 0.3299 | 0.9020 | 0.9301 | 0.3250 | 0.9020 | 0.9301 | 0.3180 |
| 5k | 10% | Cov. | 0.9018 | 0.9301 | 0.9055 | 0.9049 | 0.9308 | 0.9164 | 0.9020 | 0.9301 | 0.9149 | 0.9020 | 0.9301 | 0.9172 |
| 10k | 10% | Cov. | 0.9018 | 0.9301 | 0.9348 | 0.9020 | 0.9301 | 0.9312 | 0.9020 | 0.9301 | 0.9305 | 0.9020 | 0.9301 | 0.9308 |
| 1k | 0.1% | Pow. | 0.9073 | 0.9327 | 0.7413 | 0.9056 | 0.9307 | 0.7326 | 0.9073 | 0.9327 | 0.7417 | 0.9073 | 0.9327 | 0.7457 |
| 5k | 0.1% | Pow. | 0.9073 | 0.9327 | 0.9087 | 0.9073 | 0.9327 | 0.9120 | 0.9073 | 0.9327 | 0.9093 | 0.9073 | 0.9327 | 0.9143 |
| 10k | 0.1% | Pow. | 0.9073 | 0.9327 | 0.9270 | 0.9073 | 0.9327 | 0.9287 | 0.9073 | 0.9327 | 0.9280 | 0.9073 | 0.9327 | 0.9317 |
| 1k | 1% | Pow. | 0.9073 | 0.9327 | 0.7433 | 0.9073 | 0.9327 | 0.7320 | 0.9073 | 0.9327 | 0.7323 | 0.9073 | 0.9327 | 0.7340 |
| 5k | 1% | Pow. | 0.9093 | 0.9341 | 0.9163 | 0.9073 | 0.9327 | 0.9157 | 0.9073 | 0.9327 | 0.9140 | 0.9073 | 0.9327 | 0.9143 |
| 10k | 1% | Pow. | 0.9073 | 0.9327 | 0.9297 | 0.9073 | 0.9327 | 0.9300 | 0.9073 | 0.9327 | 0.9270 | 0.9073 | 0.9327 | 0.9297 |
| 1k | 10% | Pow. | 0.9073 | 0.9327 | 0.7257 | 0.9229 | 0.9442 | 0.7288 | 0.9073 | 0.9327 | 0.7173 | 0.9073 | 0.9327 | 0.7020 |
| 5k | 10% | Pow. | 0.9073 | 0.9327 | 0.9000 | 0.9111 | 0.9326 | 0.9100 | 0.9073 | 0.9327 | 0.9097 | 0.9073 | 0.9327 | 0.9110 |
| 10k | 10% | Pow. | 0.9073 | 0.9327 | 0.9303 | 0.9073 | 0.9327 | 0.9263 | 0.9073 | 0.9327 | 0.9257 | 0.9073 | 0.9327 | 0.9277 |

